# Neurocomputational principles underlying the number sense

**DOI:** 10.1101/2022.06.01.494401

**Authors:** Joonkoo Park, David E. Huber

## Abstract

Many species of animals exhibit an intuitive sense of number, suggesting a fundamental neural mechanism for representing numerosity in a visual scene. Recent empirical studies demonstrate that early feedforward visual responses are sensitive to numerosity of a dot array but not to continuous dimensions orthogonal to numerosity, such as size and spacing of the dots. However, the mechanisms that extract numerosity are unknown. Here we identified the core neurocomputational principles underlying these effects: (1) center-surround contrast filters; (2) at different spatial scales; with (3) divisive normalization across network units. In an untrained computational model, these principles eliminated sensitivity to size and spacing, making numerosity the main determinant of the neuronal response magnitude. Moreover, a model implementation of these principles explained both well-known and relatively novel illusions of numerosity perception across space and time. This supports the conclusion that the neural structures and feedforward processes that encode numerosity naturally produce visual illusions of numerosity. Together, these results identify a set of neurocomputational properties that gives rise to the ubiquity of the number sense in the animal kingdom.

## Introduction

Humans have an intuitive sense of number that allows numerosity estimation without counting^1^. The prevalence of number sense across phylogeny and ontogeny^2^ suggests common neural mechanisms that allow the extraction of numerosity information from a visual scene. While earlier empirical work highlighted the parietal cortex for numerosity representation^3^, growing evidence suggests that numerosity is processed at a much earlier stage. A recent study, using high-temporal resolution electroencephalography together with a novel stimulus design, demonstrated that early visual cortical activity is uniquely sensitive to the numerosity of a dot array in the absence of any behavioral response, but not to non-numerical dimensions that are orthogonal to numerosity (i.e., size and spacing, **Fig. 1A**)^4^ Subsequent behavioral and neural studies showed that this early cortical sensitivity to numerosity indicates feedforward activity in visual areas V1, V2 and V3^5–7^ These results suggest that numerosity is a basic currency of perceived magnitude early in the visual stream.

**Figure 1.**
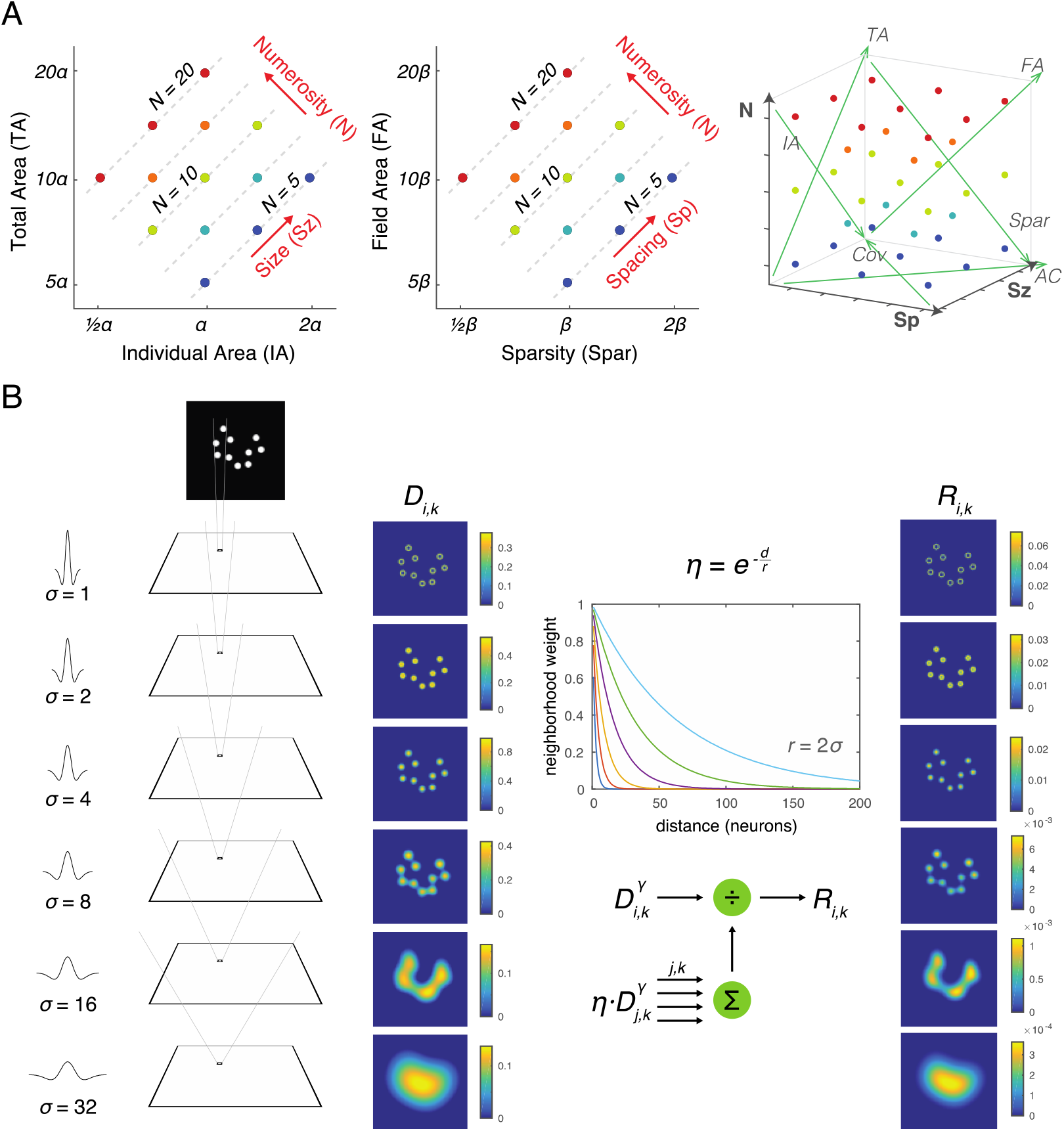
Stimulus design and computational methods. A. Properties of magnitude dimensions represented in three orthogonal axes defined by log-scaled number (N), size (Sz), and spacing (Sp) (Table 1). B. Schematic illustration of the computational process from a dot-array image to the driving input (i.e., the model without divisive normalization), D, of the simulated neurons, versus the normalized response (i.e., the model with divisive normalization), R. A bitmap image of a dot array was fed into a convolutional layer with DoG filters in six different sizes (Eq. 1). The resulting values, after half wave rectification, represented the driving input. Neighborhood weight, defined by η, was multiplied by the driving input across all the neurons across all the filter sizes, the summation of which served as the normalization factor (see Eq. 2 & 3). This illustration of η is showing the case where r is defined by twice the size of the sigma for the DoG kernel.

Nevertheless, it is unclear how feedforward neural activity creates a representation of numerosity within these brain regions. Specifically, the view of numerosity as a *discrete number* of items seems incompatible with the primary modes of information processing in the brain, such as firing rates and population codes, which are *continuous.* Indeed, some authors assume that continuous non-numerical magnitude information is encoded first and integrated to produce the representation of numerosity^8–10^. In contradiction, however, recent empirical studies demonstrate that the magnitude of visual cortical activity is most sensitive to number and is relatively insensitive to other continuous dimensions such as size and spacing of a dot array^11–14^

What explains this insensitivity to spacing and size effects, despite robust sensitivity to number? Previous computational modeling studies offer some hints to this question. The earliest computational model of numerosity perception by Dehaene and Changeux^15^ assumes that the response to each dot in an array is normalized across dot size via a convolution layer in which the filter that matches the dot size dominates the neuronal activity as a result of a winner-take-all process. At the time this model was proposed, it was not known that numerosity is encoded in early visual processing. As applied to early vision, this strong winner-take-all mechanism is implausible, as the model would suggest that visual cortex only knows that dots exist, without knowing the size or the location of the dots. Also, this model does not allow *any* effect of size or spacing, which is inconsistent with empirical data^16^.

Recently, several deep-network-based models have been applied to numerosity perception^17–21^. Stoianov and Zorzi^20^ developed a hierarchical generative model of the sensory input (images of object arrays) and demonstrated that after learning to generate its own sensory input, some units in the hidden layer were sensitive to numerosity irrespective of total area while other units were sensitive to total area irrespective of numerosity. This suggests an unsupervised learning mechanism for efficient coding of the sensory data that can extract statistical regularities of the input images. The authors provided some suggestions as to the specific neurocomputational principle(s) underlying the success of this model. For example, the first hidden layer developed center-surround representations of different sizes and the second layer developed a pattern of inhibitory connections to units in the first layer that encoded cumulative area. However, the development of center-surround detectors based on unsupervised learning is a common observation^22^, indicating that such results are not unique to displays of dot arrays, and are instead a natural byproduct of learning in the visual system. In a more recent study, Kim and colleagues^21^ found that sensitivity and selectivity to numerosity was well captured in a completely untrained convolutional neural network (AlexNet)^23^, suggesting that a repeated process of convolution and pooling is capable of normalizing continuous dimensions and extracting numerosity information as a statistical regularity of an image. However, these are “black box” models, and it is not always clear *how* these models work; these models contain many mechanisms, and it is not clear which mechanisms are crucial for producing numerosity-sensitive units.

Rather than applying a complex multilayer learning model, we distill the neurocomputational principles that enable the visual system to be sensitive to numerosity while remaining insensitive to non-numerical visual features. Consistent with prior work, we hypothesize that center-surround contrast filters at different spatial scales play an important role in numerosity perception. In contrast to prior proposals, we hypothesize that divisive normalization^24^, in which the response of each neuron reflects its driving input divided by the summation of responses from anatomically surrounding neurons, plays a key role in numerosity perception. In the case of early vision, the surround is spatially determined in terms of retinotopic positions. Divisive normalization is known to exist throughout the cortex, reflecting the shunting inhibition of inhibitory interneurons that limit neural activation within a patch of cortex^24^ Critically, unlike connections between layers, such as with the pooling layers of AlexNet, divisive normalization occurs within a layer (e.g., between center-surround units) through recurrent activation. Furthermore, a wealth of evidence indicates that divisive normalization is ubiquitous across species and brain systems and hence thought to be a fundamental computation of many neural circuits. Thus, any theory of numerosity perception would be remiss, *not* to include the effect of divisive normalization.

To determine the contribution of divisive normalization to numerosity encoding, we implemented an untrained neural network with versus without divisive normalization as applied to center-surround filters at different spatial scales (e.g., as in V1) (**Fig. 1B**). The output simulates the summation of synchronized postsynaptic activity of a large population of neurons at a pre-decisional stage, consistent with previous work^4,6^. Our results show that (1) hierarchically organized multiple center-surround filters of varying size make the network insensitive to spacing and that (2) divisive normalization implemented across network units makes the network additionally insensitive to size. Divisive normalization not only occurs over space but also over time^25^. Thus, we additionally implemented temporal divisive normalization to test if it explains contextual effects of numerosity perception^16,26^.

## Results

### Center-surround convolution captures total pixel intensities and eliminates the effect of spacing

Images of dot arrays that varied systematically across number, size, and spacing (see Materials and Methods) were fed into a convolutional layer with difference-of-Gaussians (DoG) filters in six different sizes. The driving input, *D,* for each filter was the convolution of a DoG with the display image, or a weighted sum of local pixel intensities (**Fig. 1B**). The summed driving input in each filter size showed different effects as a function of number, size, and spacing (**Fig. 2A**), but when the driving input was summed across all filter sizes it was most strongly modulated by both number and size equally but not by spacing (**Fig. 2B**), suggesting that the neural activity tracks total area *(TA;* see **Table 1**; **Fig. S1**). The effect of spacing existed in the fourth and sixth largest filter sizes, largely indicating effects of field area and density, respectively (**Fig. 2A**); however, the effects in these two filter sizes were in opposite directions, which made the overall effect very small. These results illustrate that having multiple filter sizes is key to normalizing the spacing dimension. In sum, the driving input of the convolutional layer captured total pixel intensity of the image regardless of the number or spatial configuration of dots.

**Figure 2.**
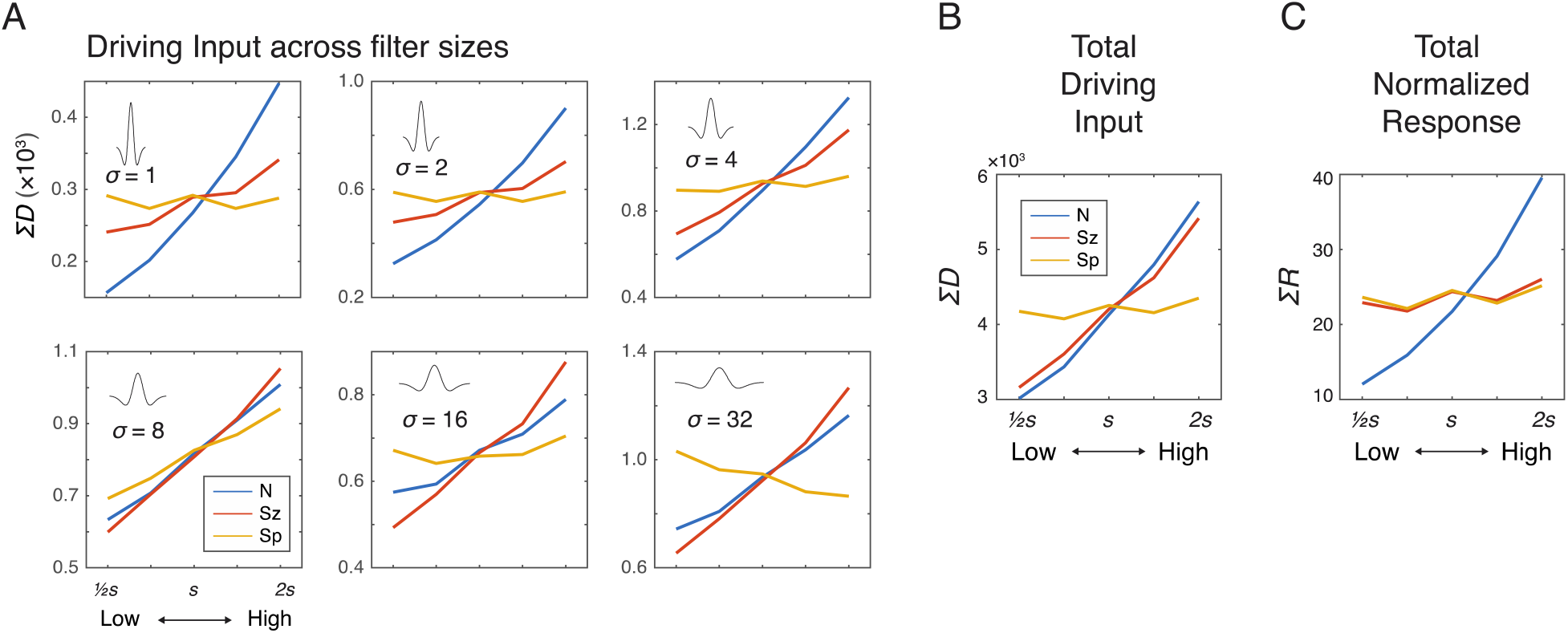
Simulation results showing the effects of N, Sz, and Sp on the driving input and normalized response of the network units. A. Summed driving input (ΣD) separately for each of the six filter sizes as a function of N, Sz, and Sp (see Methods for the specific values of s). B. ΣD across all filters is modulated by both number and size but not by spacing. C. The summed normalized response showed a near elimination of the Sz effect leaving only the effect of N. The results were simulated using r = 2σ and γ = 2, but effects of Sz and Sp were negligible across all the tested model parameters (Fig. S2). The value s on the horizontal axis indicates a median value for each dimension (see Materials and Methods).

**Table 1.** Mathematical relationship between various magnitude dimensions.

| Dimension | As a function of $n$, $r_d$, $r_f$ | As a function of $N$, $S_z$, $Sp$ |
| --- | --- | --- |
| Individual area (IA) | $\pi r_d^2$ | $\log(IA) = \frac{1}{2}\log(S_z) - \frac{1}{2}\log(N)$ |
| Total area (TA) | $n \times \pi r_d^2$ | $\log(TA) = \frac{1}{2}\log(S_z) + \frac{1}{2}\log(N)$ |
| Field area (FA) | $\pi r_f^2$ | $\log(FA) = \frac{1}{2}\log(Sp) + \frac{1}{2}\log(N)$ |
| Sparsity (Spar) | $\pi r_f^2 / n$ | $\log(Spar) = \frac{1}{2}\log(Sp) - \frac{1}{2}\log(N)$ |
| Individual perimeter (IP) | $2\pi r_d$ | $\log(IP) = \log(2\sqrt{\pi}) + \frac{1}{4}\log(S_z) - \frac{1}{4}\log(N)$ |
| Total perimeter (TP) | $n \times 2\pi r_d$ | $\log(TP) = \log(2\sqrt{\pi}) + \frac{1}{4}\log(S_z) + \frac{3}{4}\log(N)$ |
| Coverage (Cov) | $n \times r_d^2 / r_f^2$ | $\log(Cov) = \frac{1}{2}\log(S_z) - \frac{1}{2}\log(Sp)$ |
| Closeness (Close) | $\pi^2 \times r_d^2 \times r_f^2$ | $\log(Close) = \frac{1}{2}\log(S_z) + \frac{1}{2}\log(Sp)$ |

Note: $n$ = number; $r_d$ = radius of individual dot; $r_f$ = radius of the invisible circular field in which the dots are drawn.

### Divisive normalization nearly eliminates the effect of size

We next added divisive normalization to the center-surround model, with different parameter values (neighborhood size and amplification factor) to determine the conditions under which divisive normalization might reduce or eliminate the effect of size and whether it might alter the absence of spacing effects in the driving input. Driving input was normalized by the normalization factor defined by a weighted summation of neighboring neurons and filter sizes (**Eq. 2**). The summed normalized responses, Σ*R*, were strongly modulated by number but much less so, if any, by size and spacing (**Fig. 2C**). The pattern of results was largely consistent across different parameter values for neighborhood size (*r*) and amplification factor (*γ*) of the normalization model (**Fig. S2**); therefore, we chose moderate values of *r* (=2) and *γ* (=2) for subsequent simulations. A regression model with summed normalized responses as the dependent measure and the three orthogonal dimensions (*N, Sz, Sp*) as the independent variables revealed a much larger coefficient estimate for *N*(*b* = 13.68) than for *Sz* (*b* = 1.541) and for *Sp* (*b* = 0.7809). In sum, a modest degree of divisive normalization eliminates the effect of size and, at the same time, does not alter the absence of spacing effects.

### Divisive normalization across space explains various visual illusions

Next, we considered if the center-surround model with divisive normalization also explains some of the most well-known visual illusions of numerosity perception. If so, this would support the hypothesis that these visual illusions reflect early visual processing at the level of numerosity encoding, without requiring any downstream processing. In other words, early vision may be the root cause of *both* numerosity encoding and numerosity visual illusions.

Empirical studies have long shown that irregularly spaced arrays (compared with regularly spaced arrays), arrays with spatially grouped items (compared with ungrouped items), and arrays with dots that are pairwise connected (compared with unconnected arrays) are all underestimated^27–31^. All three of these illusions were captured by the inclusion of divisive normalization (**Fig. 3A-C**), whereas in the absence of divisive normalization, there was either no effect or an effect in the opposite direction (**Fig. S3**). The underestimation effects in the normalized response can be explained by greater normalization when neurons with overlapping normalization neighborhoods are activated, with this greater overlap occurring in subregions of the images for irregular, grouped, or connected (lines) dots. This explanation is functionally similar to one provided by the “occupancy model”^32^, but our results demonstrate that these effects emerge naturally within early visual processing.

**Figure 3.**
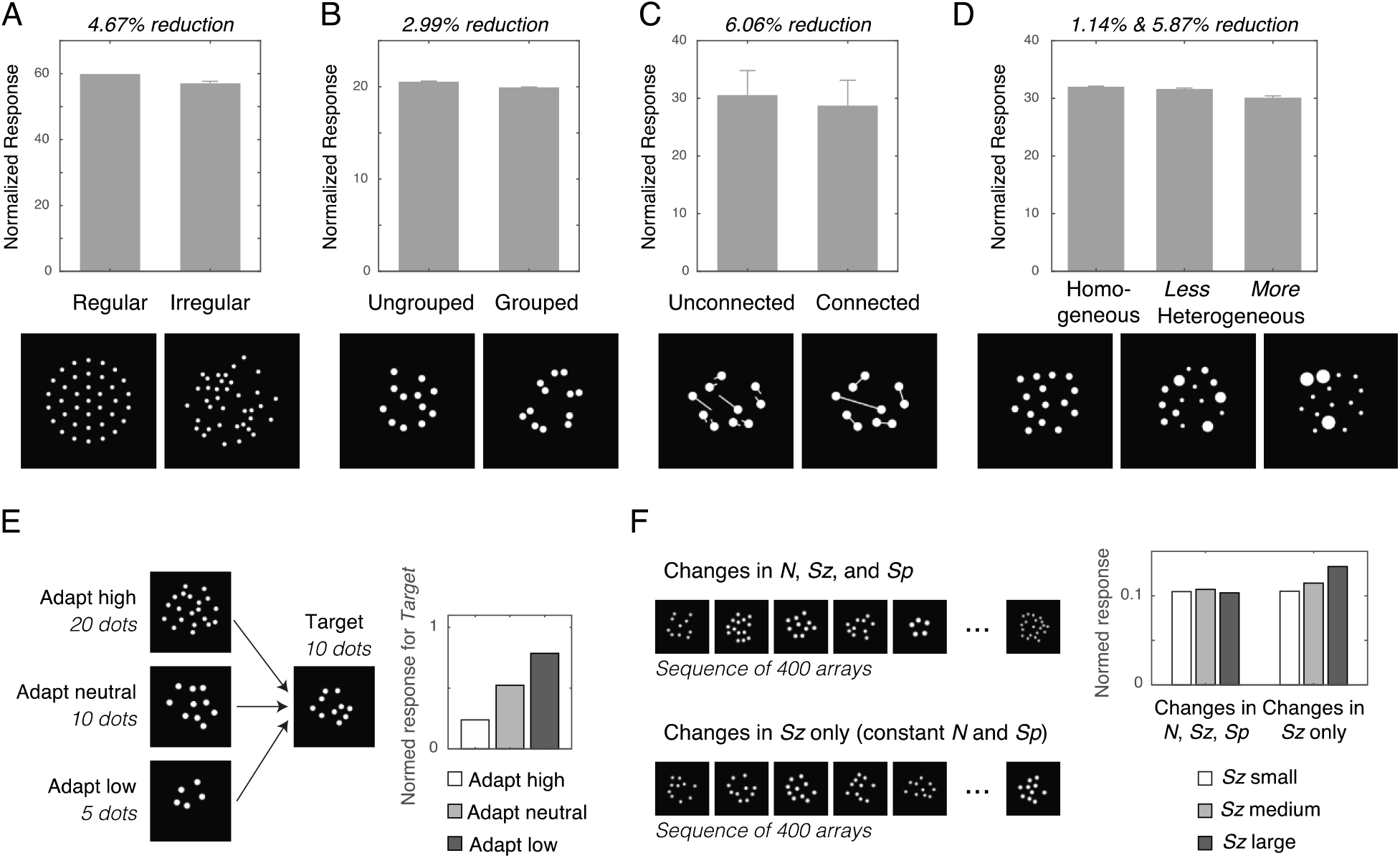
Simulation of numerosity illusions. Normalized response of the network units influenced by the (A) regularity, (B) grouping, (C), connectedness, and (D) heterogeneity of dot arrays, as well as by (E) adaptation and (F) context. Error bars represent one standard deviation of the normalized response across simulations. In the case of temporal normalization effects (E and F), the results were simulated with ω = 8 and δ = 2, but the patterns held across all tested model parameter values.

A relatively understudied visual illusion is the effect of heterogeneity of dot size on numerosity perception. A recent behavioral study demonstrated that the point of subjective equality was about 5.5% lower in dot arrays with heterogenous sizes compared with dot arrays with homogeneous sizes^33^. Consistent with this behavioral phenomenon, our simulations revealed that greater heterogeneity leads to greater underestimation (**Fig. 3D**). This occurs because the summed normalized response of a single dot saturates as dot area increases (**Fig. S4**), which interacts with the heterogeneity of the dot array. As heterogeneity is manipulated by making some dots larger and other dots smaller while keeping total area and numerosity constant, this saturating effect makes the overall normalized response smaller as a greater number of dots deviates from the average size (the gains from making some dots larger is not as great as the losses from making some dots smaller). As in the case of other illusions, the same analysis in the absence of divisive normalization fails to produce this illusion (**Fig. S3**).

### Divisive normalization across time explains numerosity adaptation and context effects

One of the most well-known visual illusions in numerosity perception is the adaptation effect^26^. We reasoned that numerosity adaptation might reflect divisive normalization across time, similar to adaptation with light or odor^24^, which shifts the response curve and produces a contrast aftereffect. Closely related to temporal adaptation, the recently discovered temporal contextual effect of numerosity perception is an amplified neural response to changes in one dimension (e.g., changes in dot size) when observers experience a trial sequence with only changes in that dimension^16^. Therefore, we also applied the model with temporal normalization to the context effect.

We modeled temporal divisive normalization for readout neurons that are driven by the sum of the normalized responses across all units, Σ*R*. This summed total response (now referred to as *M*) was temporally normalized (*M*\*) by the recency weighted average of the driving input (**Eq. 4**). Temporal normalization shifts the sigmoid response curve horizontally along the dimension of *M* to maximize the sensitivity of *M*\* based on the recent history of stimulation. Provided that the constant in the denominator is approximately equal to the current trial’s response, the results of spatial normalization reported above would not change by also introducing temporal normalization. Temporal normalization was assessed for cases of a target array of 10 dots after observing an array of 5 dots, 10 dots, or 20 dots (**Fig. 3E**). Similar to behavioral results^34^, the target of 10 dots was underestimated when the adaptor was more numerous than the target and was overestimated when the adaptor was less numerous than the target. These results confirm that divisive normalization across time naturally produces numerosity adaptation.

Using the same model of temporal normalization (**Eq. 4**), we tested if it can also explain longer-sequence context effects. Studies show that the effect of size is negligible in the context of a trial sequence that varies size, spacing, and number^4^, but that the effect of size becomes apparent when number and spacing are held constant while varying only size^16^. We simulated each of these contexts: The model saw a total of 400 dot arrays that varied across number, size, and spacing or else it saw 400 dot arrays that differed only in size (**Fig. 3F**). In the context where all dimensions varied, the three levels of *Sz* had no linear association with *M*\*; the 95 percentile confidence interval of the ordinary-least-square linear slope of *M*\* as a function of *Sz* was [-0.0411, 0.0293], which includes 0. In contrast, in the context where only size varied, *M\** was positively correlated with *Sz*; slope confidence interval of [0.00655, 0.00725], which excludes 0. This phenomenon can be explained by the adaptive shifting of the sigmoid response curve across trials. In the former case, because recent trials are often of larger or smaller total response as compared to the current trial, the normalization for the current trial is more often pushed to the nonlinear parts of the normalization curve (e.g., closer to ceiling and floor effects). Thus, the temporally normalized response is relatively insensitive to the small effect of size (keeping in mind that the effect of size is made small by spatial divisive normalization). In contrast, when only size varies across trials, the total response of recent trials is more likely to be well-matched to the total response of the current trial. As a result, the small effect of size is magnified in light of this temporal stability.

## Discussion

Despite the ubiquity of number sense across animal species, it was previously unclear how unadulterated perceptual responses produce the full variety of numerosity perception effects. Recent empirical studies demonstrate that feedforward neural activity in early visual areas is uniquely sensitive to the numerosity but much less so, if any, to the dimension of size and spacing, which are continuous non-numerical dimensions that are orthogonal to numerosity. Despite recent advances showing that numerosity information *can* be extracted from a deep neural network^17,20,21^, precisely *how* early visual areas normalize the effects of size and spacing was unclear.

The current study identified the key neurocomputational principles involved in this process. First, the implementation of hierarchically organized multiple sizes of center-surround filters effectively normalizes spacing owing to offsetting factors (**Fig. 4A**). On the one hand, relatively smaller filters that roughly match or are slightly bigger than each dot produce a greater response when the dots are farther apart because their off-surround receptive fields do not overlap. On the other hand, relatively larger filters that cover most of the array produce a greater response when the dots are closer together because stimulation at the center of the on-surround receptive fields is maximized. When summing these opposing effects, which occur at different center-surround filter sizes, the overall neural activity is relatively invariant to spacing. Second, the implementation of divisive normalization reduces the effect of size by reducing activity at larger filter sizes that have overlapping normalization neighborhoods (**Fig. 4B**). More specifically, increase in size produces greater overall unnormalized activity because more filters (e.g., both larger and smaller) are involved in responding to larger dots whereas only smaller filters respond to small dots (**Fig. 2B**). However, normalization dampens this increase. Critically, divisive normalization is a within-layer effect, reflecting recurrent inhibition between center-surround filters owing to inhibitory interneurons. Thus, the effect of dot size is eliminated in early visual responses. In sum, contrast filters at different spatial scales and divisive normalization naturally increases sensitivity to the number of items in a visual scene. Because these neurocomputational principles are commonly found in visual animals, this suggests that numerosity is a natural property of the visual system.

**Figure 4.**
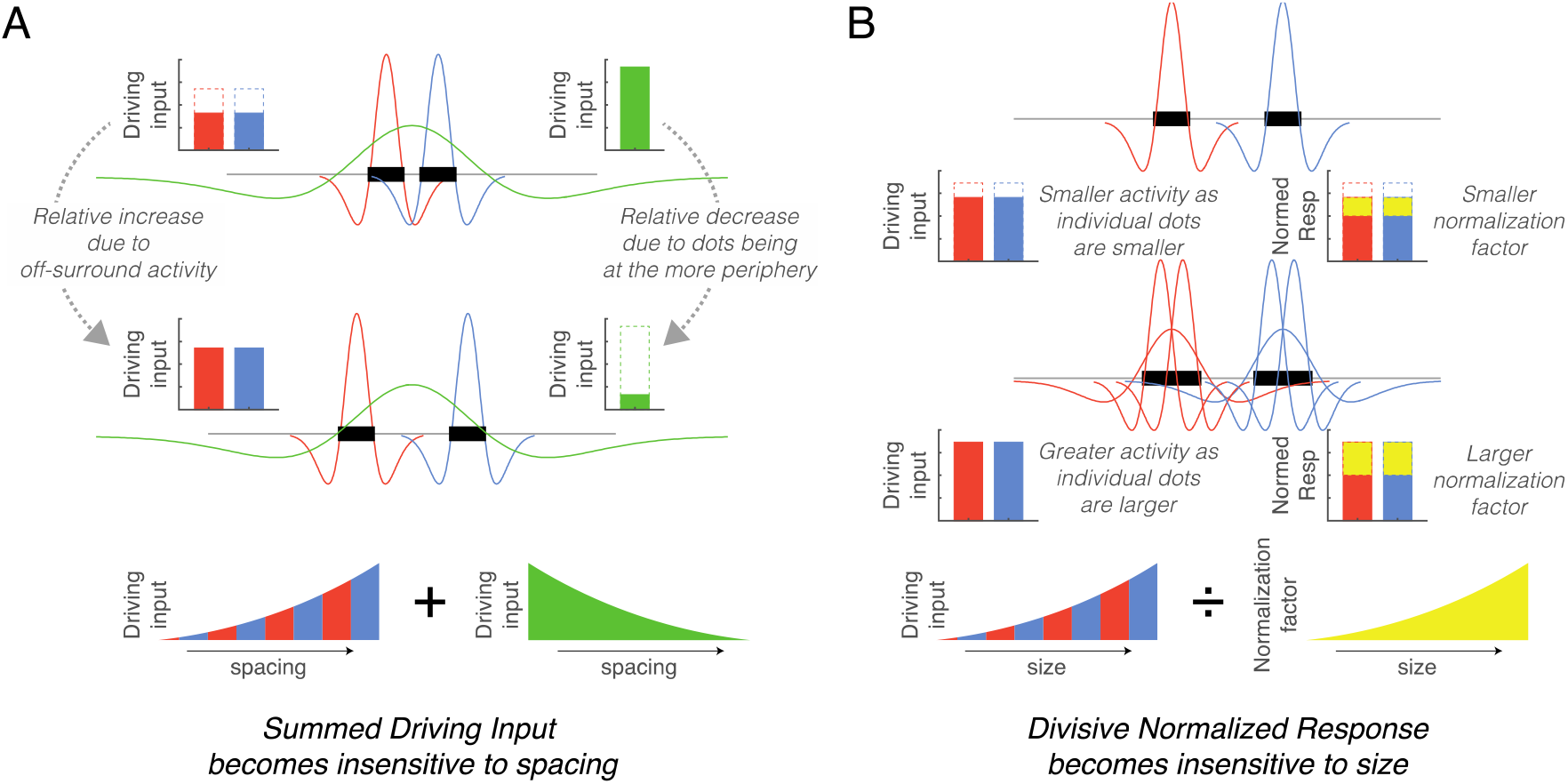
Simplified schematics explaining the mechanisms underlying the normalization of size and spacing. A. As spacing increases (from top to middle row) the response of small size center-surround filters increases (red and blue) whereas the response of large size center-surround filters decreases (green), with these effects counteracting each other in the total response. B. As dot size increases (from top to middle row), more filters are involved in responding to the dots thereby increasing the unnormalized response (red and blue), but this results in a greater overlap in the neighborhoods and increases the normalization factor (yellow). These counteracting effects eliminate the size effect.

A key result from the current model is that the summed normalized output of the neuronal activity is sensitive to numerosity but shows little variation with size and spacing. This pattern is consistent with electroencephalography studies finding similar results for the summed response of V1, V2, and V3 in the absence of any behavioral judgment^6,7^ However, this pattern is different than the behavior of prior deep neural network-based models of numerosity perception, which revealed many units in the deep layers that were sensitive to non-numerical dimensions, along with a few that were numerosity sensitive (or selective). Although the few units that were selective to numerosity could explain behavior, the abundance of simulated neurons sensitive to non-numerical dimensions is inconsistent with population-level neural activity, which fails to show sensitivity to these non-numerical dimensions in early visual cortex^11–14^ A key difference between the current model and previous computational models is the inclusion of divisive normalization in the center-surround convolution layer. Unlike prior models, this eliminated the effect of size in the early visual response, without requiring subsequent pooling layers^17–21^ or a decision making process that compares high versus low spatial frequency responses^35^. In summary, our results suggest that (1) center-surround filters, (2) of varying filter sizes, with (3) divisive normalization, naturally reduce the effects of size and spacing while maintaining sensitivity to number without requiring second stage processing.

Casting doubt on our conclusion that numerosity arises from center-surround filters and divisive normalization, a recent fMRI study finds that visual cortical activity evoked by dot arrays follows aggregate Fourier power more closely than numerosity^14^ However, there are two caveats that make it unclear if this study contradicts our claim. First, Fourier power is based on the analysis of spatially unbounded sine waves that has little biological plausibility (unlike center-surround or Gabor filters, which are spatially limited). Second, more critically, aggregate Fourier power in that study was quantified only up to a certain harmonic of the image. In other words, the Fourier metric used in the analysis required a sort of homunculus that tailored low-pass frequency filtering based on dot size. Hence, it is not surprising that aggregate Fourier power up to the frequency defined by dot size in each image follows the number of dots in that image. If all frequencies had been included, smaller dots would have produced an overall larger Fourier power than was assumed in the analysis, which would likely change the conclusions. In addition, this dot-size truncated version of Fourier power failed to simulate the connectedness illusion whereas the center-surround model with divisive normalization correctly produced a decreases response (**Fig. 3C**), consistent with behavior and neuroimaging^5,7,29^

Our conclusions are primarily in terms of the qualitative effects of center-surround filtering and divisive normalization, which collectively produce selective sensitivity to numerosity. However, specific quantitative predictions will change depending on specific model assumptions. For instance, our simulations assumed a distribution of filter sizes that ranged from much smaller to much larger than the presented dots, although this assumption might be incorrect when considering dots in the periphery where receptive field sizes are larger^36,37^ or if the dots are crowded and hard to individuate^38^. In fact, both of these cases are known to exhibit different behavioral characteristics from the perception of centrally presented, uncrowded dot arrays. Our simulation also assumed an equal number of small and large center-surround filters although in reality there are likely fewer large filters. This assumption was made out of computational convenience, although we note that similar results would emerge with an unequal distribution of filters if the divisive normalization amplification factor scaled with filter size (e.g., if the larger number of small filters more strongly inhibited each other) or if the neighborhood size of divisive normalization scaled with filter size in a nonlinear manner.

A primary feature of early visual cortex regions such as V1 and V2 is oriented line filters (i.e., simple cells). The fact that center-surround DoG filters in a feedforward network were sufficient for explaining numerosity perception suggests that neural activity at even earlier stages of the visual pathway, such as the retina or the lateral geniculate nuclei, may be capable of representing numerosity. Consistent with this idea, recent studies show evidence for numerosity processing in the subcortical structure following the monocular visual pathway^39^ and in the nervous structure that gives rise to the modulation of the pupillary response^40^.

The success of this model does not necessarily imply that neuronal responses in early visual regions directly determine behavioral responses^7^. Prior to behavior, there are many downstream processing steps that incorporate other sources of information, such response bias and decisional uncertainty. Instead, these results, together with previous electrophysiology results, suggest that normalized response magnitude in early visual regions may be the basic currency from which numerosity judgments are made. Future work should explore the link between the neuronal response layer in the current model and various behavioral judgments. For instance, if decisional uncertainty is modeled by assuming a constant level of decisional noise, regardless of numerosity, then the model will naturally produce Weber’s scaling law of just noticeable differences considering that the normalized response follows a log-linear pattern as a function of numerosity (see **Fig. 2C**). More complex decisional assumptions could be introduced in an attempt to model the effects of task instructions that are known to bias decisions on magnitude judgment^41,42^. More assumptions about top-down semantic influences may also explain recent coherence illusion results in orientation or color^43,44^, for instance if observers are drawn to focus on a particular feature of the stimulus when comparing two dot arrays.

Another line of possible future work concerns divisive normalization in higher cortical levels involving neurons with more complex receptive fields. While the current normalization model with center-surround filters successfully explained visual illusions caused by regularity, grouping, connectedness, and heterogeneity, other numerosity phenomena such as statistical pairing and topological invariants^30,45^ may require the action of neurons with receptive fields that are more complex than center-surround filters. Inclusion of more complex receptive fields may improve the explanatory power of the model in some cases. For instance, although the normalized response was reduced by 6% for arrays of connected dots in our simulation, the equivalent reduction in behavior is about 20%, suggesting that this is not a full account of the connectedness effect. However, oriented line filters (Gabors capturing higher spatial frequency) would better capture the shape of the connected dots (e.g., barbell like figures), which should enhance the connectedness effect. This would help explain recent studies suggesting that the connectedness effect reflects two sweeps of feedforward response (low spatial frequency at 100 ms and high spatial frequency at 150 ms) in visual cortex^5,7^

In conclusion, our results indicate that divisive normalization in a single convolutional layer with hierarchically organized center-surround filters naturally enhances sensitivity to the discrete number of items in a visual scene by reducing the effects of size and spacing, consistent with recent empirical studies demonstrating direct and rapid encoding of numerosity^4^ This account predicts that various well-known numerosity illusions across space and time arise naturally within the same neural responses that encode numerosity, rather than reflecting later stage processes. These results identify the key neurocomputational principles underlying the ubiquity of the number sense in the animal kingdom.

## Methods

### Stimulus sets

#### Dot arrays spanning across number, size, and spacing

Inputs to the neural network were visual stimuli of white dot arrays on a black background (200 × 200 pixels). Dots were homogeneous in size within an array and were drawn within an invisible circular field. Any two dots in an array were at least a diameter apart from edge to edge. The number of dots in an array is referred to as *n*, the radius of each dot is referred to as *r_d_*, and the radius of the invisible circular field is referred to as *r_f_*. **Table 1** provides mathematical definitions of other non-numerical dimensions based on these terms.

Following the previously developed framework for systematic dot array construction^4,46^, stimulus parameters of the dot arrays were distributed systematically within a parameter space defined by three orthogonal dimensions: log-scaled dimensions of number (*N*), size (*Sz*), and spacing (*Sp*) (**Fig. 1A**). *N* simply represents the number of dots. *Sz* is defined as the dimension that varies with individual area (*IA*) while holding *N* constant, hence simultaneously varying in total area (*TA*). *Sp* is defined as the dimension that varies with sparsity *(Spar)* while holding *N* constant, hence simultaneously varying in field area (*FA*). Log-scaling these dimensions allows *N, Sz,* and *Sp* to be orthogonal to each other and represent all of the non-numerical dimensions of interest to be represented as a linear combination of those three dimensions (see **Table 1**). Thus, this stimulus construction framework makes is easy to visualize the stimulus parameters and analyze choice behavior or neural data using a linear statistical model. For an implementation of this framework, see the MATLAB code published in the following public repository: https://osf.io/s7xer/.

Across all the dot arrays, number (*n*) ranged between 5 to 20 dots, dot diameter (2 × *r_d_*) ranged between 9 to 18 pixels, field radius (*r_f_*) ranged between 45 to 90 pixels, all having five levels in logarithmic scale. log(*N*) ranged from 2.322 to 4.322 with the median of 3.322; log(*Sz*) ranged from 16.305 to 18.305 with the median of 17.305; log(*Sp*) ranged from 19.646 to 21.646 with the median of 20.646. This approach resulted in 35 unique points in the three-dimensional parameter space (see **Fig. 1A**). For each of the 35 unique points, a total of 100 dot arrays were randomly constructed for the simulation conducted in this study.

#### Dot arrays for testing regularity effects

The ‘regular’ dot array was constructed following the previous study that first demonstrated the regularity effect^27^ This array contained 37 dots with *r_d_* = 3 pixels, one of which at the center of the image and the rest distributed in three concentric circles with the radii of 20, 40, and 60 pixels. The ‘irregular’ arrays were constructed with the same number of and same sized dots randomly placed with *r_f_* = 72.5 pixels. This radius for the field area was empirically calculated so that the convex hull of the regular array and the mean convex hull of the irregular arrays were matched. Sixteen irregular arrays were used in the simulation.

#### Dot arrays for testing grouping effects

One set of ‘ungrouped’ dot arrays and another set of ‘grouped’ dot arrays were constructed. Both ungrouped and grouped arrays contained 12 dots, each of which with *r_d_* = 4.5 pixels. However, in the ungrouped arrays the dots were randomly dispersed, while in the grouped arrays the dots were spatially grouped in pairs. The edge-to-edge distance between the two dots in each pair was approximately equal to *r_d_*. A large number of unique dot arrays were constructed using these criteria for each of the two sets. Then, a subset of unique arrays from each set was chosen so that the convex hull of the arrays between the two sets were numerically matched. A total of 16 grouped and 16 ungrouped arrays entered the simulation.

#### Dot arrays for testing connectedness effects

One set of ‘connected’ dot arrays and another set of ‘unconnected’ dot arrays were constructed. First, a large number of dot arrays with *n* = 10, *r_d_* = 6.5 pixels, and *r_f_* = 64 pixels were created. Then, connected dot arrays were constructed by connecting the centers of two closest dots with a thin white line that was 2 pixels in width. The resulting images were visually checked, and all the images in which the lines cross or touch other lines or dots were removed from the set. Then, unconnected dot arrays were constructed from those connected dot arrays by breaking the midpoints of the interconnecting lines and rotating those broken lines about the center of each dot by ±30 degrees in either direction randomly determined. A total of 16 connected and 16 unconnected arrays entered the simulation.

#### Dot arrays for testing heterogeneity effects

Three sets of dot arrays equated in the total area (*TA*) were created. The first set of ‘homogeneous’ (or zero level of heterogeneity) dot arrays contained *n* = 15 with *r_d_* = 5 pixels within a circular field defined by *r_f_* = 75 pixels. The second set of ‘less heterogeneous’ dot arrays contained six dots with *r_d_* = 3 pixels, six dots with *r_d_* = 5 pixels, and three dots with *r_d_* = 7.5 pixels. The last set of ‘more heterogeneous’ dot arrays contained twelve dots with *r_d_* = 2.5 pixels and three dots with *r_d_* = 10 pixels. Hence, the total area (*TA*) of all the arrays were approximately identical to each other while the variability of individual area *(IA)* differed across the sets. Rounding errors due to pixelation and anti-aliasing, however, caused differences the actual cumulative intensity measure of the bitmap images. On average, the cumulative intensity values (0 being black and 1 being white in the bitmap image) were comparable between the three sets of arrays: 1209 in the homogeneous arrays, 1194 in the less heterogeneous arrays, and 1204 in the more heterogeneous arrays. Sixteen arrays in each of the three sets entered the simulation.

### Neural network model with divisive normalization

#### Convolution with DOG filters

The model consisted of a convolutional layer with difference-of-Gaussians (DoG) filters of six different sizes, that convolved input values of the aforementioned bitmap images displaying dot arrays. This architecture hence provided a structure for 200 × 200 × 6 network units (or simulated neurons) activated by images of dot arrays (**Fig. 2**). The DoG filters are formally defined as:

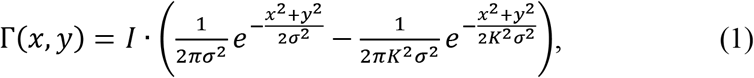

Where *I* is the input image, *σ*^2^ is the spatial variance of the narrower Gaussian, and *K* is the scaling factor between the two variances. As recommended by Marr and Hildreth^47^, *K* = 1.6 was used to achieve balanced bandwidth and sensitivity of the filters. Considering that the input values range [0 1], the DoG filters were reweighted so that the sum of the positive portion equals to 1 and the sum of the negative portion equals to −1, making the summation across all domains 0. This reweighting ensured that the response is maximized when the input matches the DoG filter regardless of filter size and that the filter produces a response of value 0 if the input is constant across a region regardless of filter size. Finally, the output of this convolution process was followed by half-wave rectification at each simulated neuron^48^, where negative responses were replaced by zero. This stipulation sets the ‘firing threshold’ of the network such that the simulated neurons would not fire if the input does not match its DoG filter.

Six different *σ* values were used (*σ_k_* = 1, 2, 4, 8, 16, 32 for filter size *k*, respectively) which together were sensitive enough to represent various visual features of the input images, from the edge of the smallest dots to the overall landscape of the entire array. The activity of each stimulated neuron, *i*, in filter size *k* following this convolution procedure is referred to as *D_i,k_*.

#### Divisive normalization

Following Carandini & Heeger^24^, the normalization model was defined as:

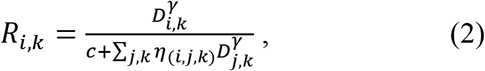

where distance similarity *η*(*ij*) is defined as:

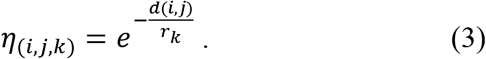

D_i_ is the driving input of neuron *i* (i.e., the output of the convolution procedure described above), *d*_(*i,j*)_ is the Euclidean distance between neuron *i* and neuron *j* in any filter size, *c* is a constant that prevents division by zero. The denominator minus this constant, which was set to 1, is referred to as the normalization factor. The parameter *r_k_*, defined for each filter size, serves to scale between local and global normalization. As *r_k_* gets larger, activities from broader set of neurons constitute the normalization factor. In our model, *r_k_* was defined as a scaling factor of *σ_k_* (e.g., *r_k_* = *σ_k_, r_k_* = 2*σ_k_*, or *r_k_* = 4*σ_k_*), so that neurons with larger filter sizes have their normalization factor computed from broader pool of neighboring neurons. The parameter *γ* determines the degree of amplification of individual inputs and serves to scale between winner take all and linear normalization. *R_i,k_* represents the normalized response of neuron *i* in filter size *k*.

#### Modelling temporal modulation of network units

Normalized responses of simulated neurons were further modeled to capture temporal modulations, with another normalization process this time working across time. First, a read out neuron was assumed that summed up the normalized responses across all the neurons, Σ*R_i,k_*. This single firing activity, now referred to as *M*, underwent the following temporal normalization process that resulted in the normalized activity *M*\*:

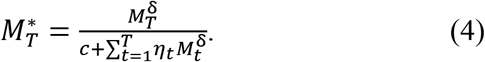

The temporal distance *η* is defined as:

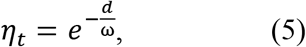

where *d* is the distance between time point *t* and *T*. As in Eq. 2 and 3, *c* is a constant that prevents division by zero, which was set to 1 for convenience. The parameter *ω* determines the amount of recent history contributing to the normalization factor, and the parameter *δ* determines the degree of amplification of *M_t_*.

The MATLAB code used to implement the model can be found in the following public repository: https://osf.io/4rwjs/ (will be made public upon acceptance).

## Acknowledgements

We thank Dr. Michele Fornaciai for inspiring discussions. This work was supported by the National Science Foundation CAREER Award BCS 1654089 to J.P.

## Author contributions

Joonkoo Park: Conceptualization, Methodology, Software, Formal analysis, Writing. David E. Huber: Methodology, Formal analysis, Writing.

## Declaration of interests

The authors declare no competing interests.

## Supplementary Figures

**Figure S1.**
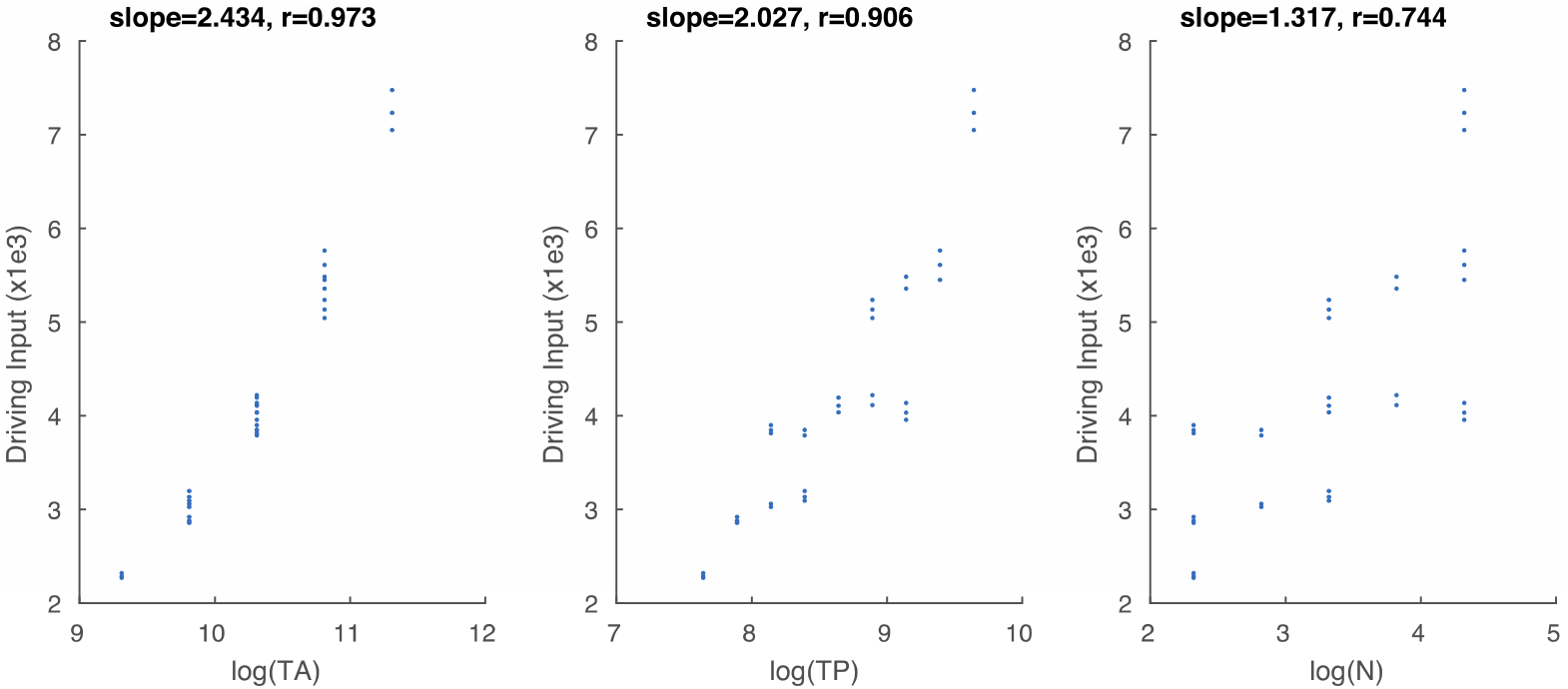
Correlation between summed driving input, Σ*D*, and log-scaled total area (TA), total perimeter (TP), and number (N).

**Figure S2.**
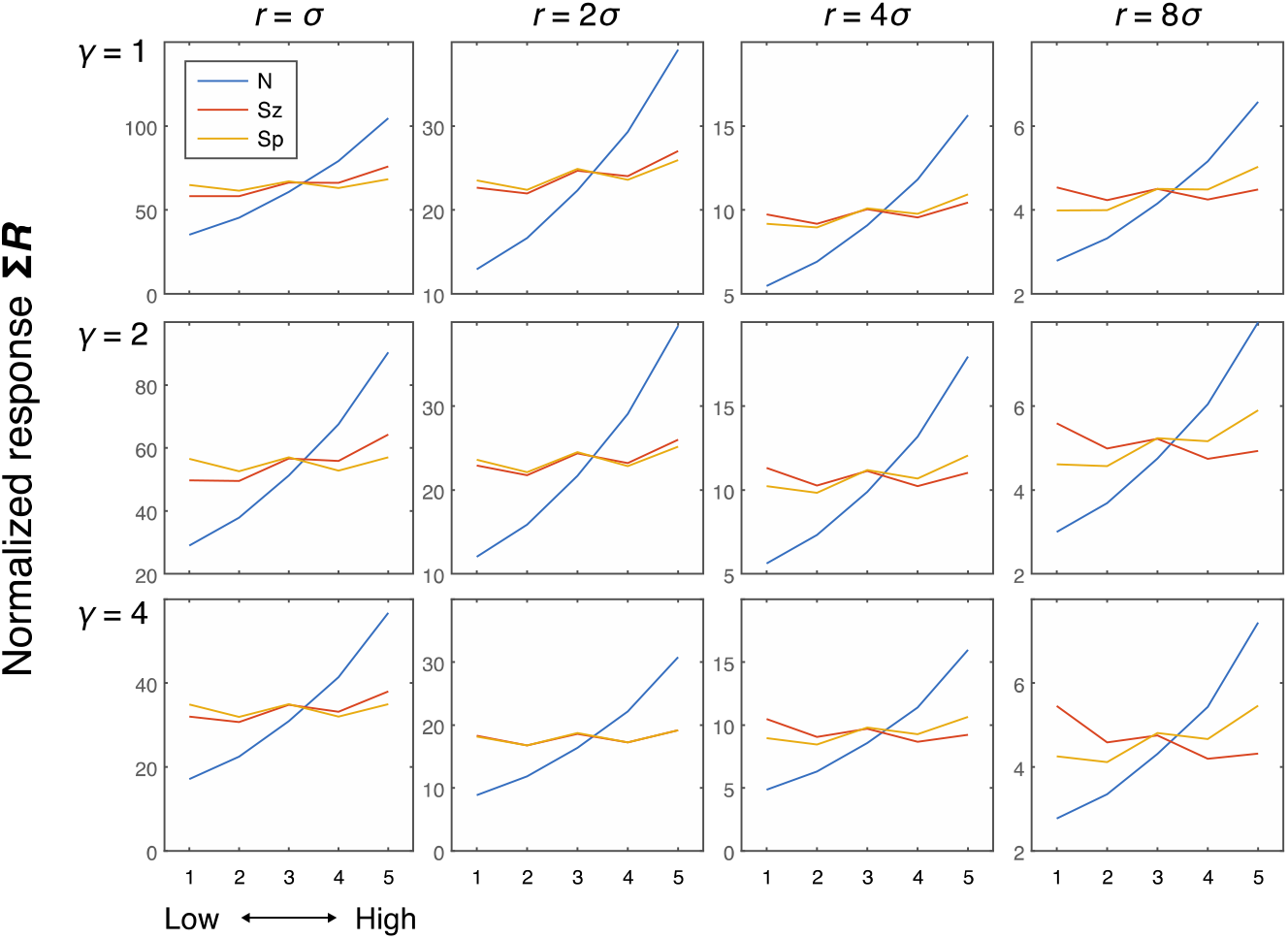
Simulation results showing the effects of number (N), size (Sz), and spacing (Sp) on the normalized response (i.e., the model with divisive normalization) of the network units as a function of neighborhood size (*r*) and amplification factor (*γ*). Greater *r* resulted in a flatter curve for the size effect, and this flattening became more pronounced as *γ* increased, with the combination of high values for both parameters producing a modest negative effect of size as well as a modest positive effect of spacing. More specifically, the combination of high *r* and *γ* values produces a winner-take-all process across large regions of the display. Greater size, in these cases, thus leads to greater normalization factor (denominator) which results in reduced normalization activity, although the extent of this normalization depends on how far away the other dots are located (e.g., less normalization with spacing). Although this is an interesting phenomenon, empirical neural and behavioral studies show a positive effect of size, if any. Hence, larger values of *r* and *γ* in this model do not seem to be plausible in the case of numerosity perception. Therefore, we chose moderate values of *r* (=2) and *γ* (=2) for subsequent simulations.

**Figure S3.**
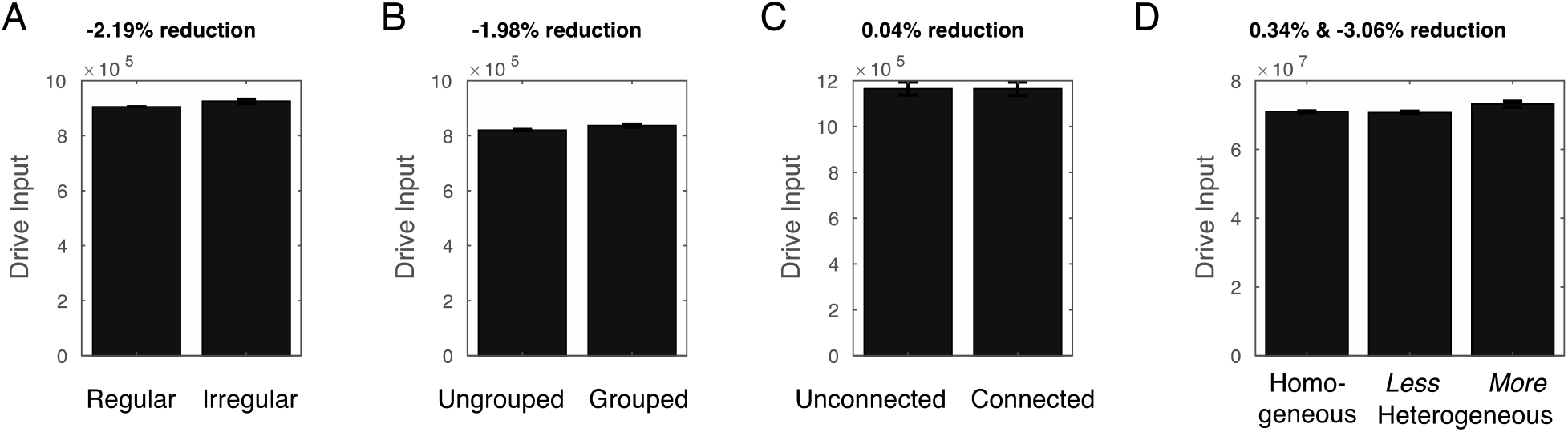
Simulation of visual illusions considering the driving input (i.e., the model without divisive normalization). No underestimation was observed in any of these cases (i.e., without divisive normalization, the model failed to explain the typically observed visual illusions). If any, irregularly spaced arrays (by 2.19%), grouped arrays (by 1.98%), and more heterogeneous arrays (by 3.06%) were overestimated based on their driving input.

**Figure S4.**
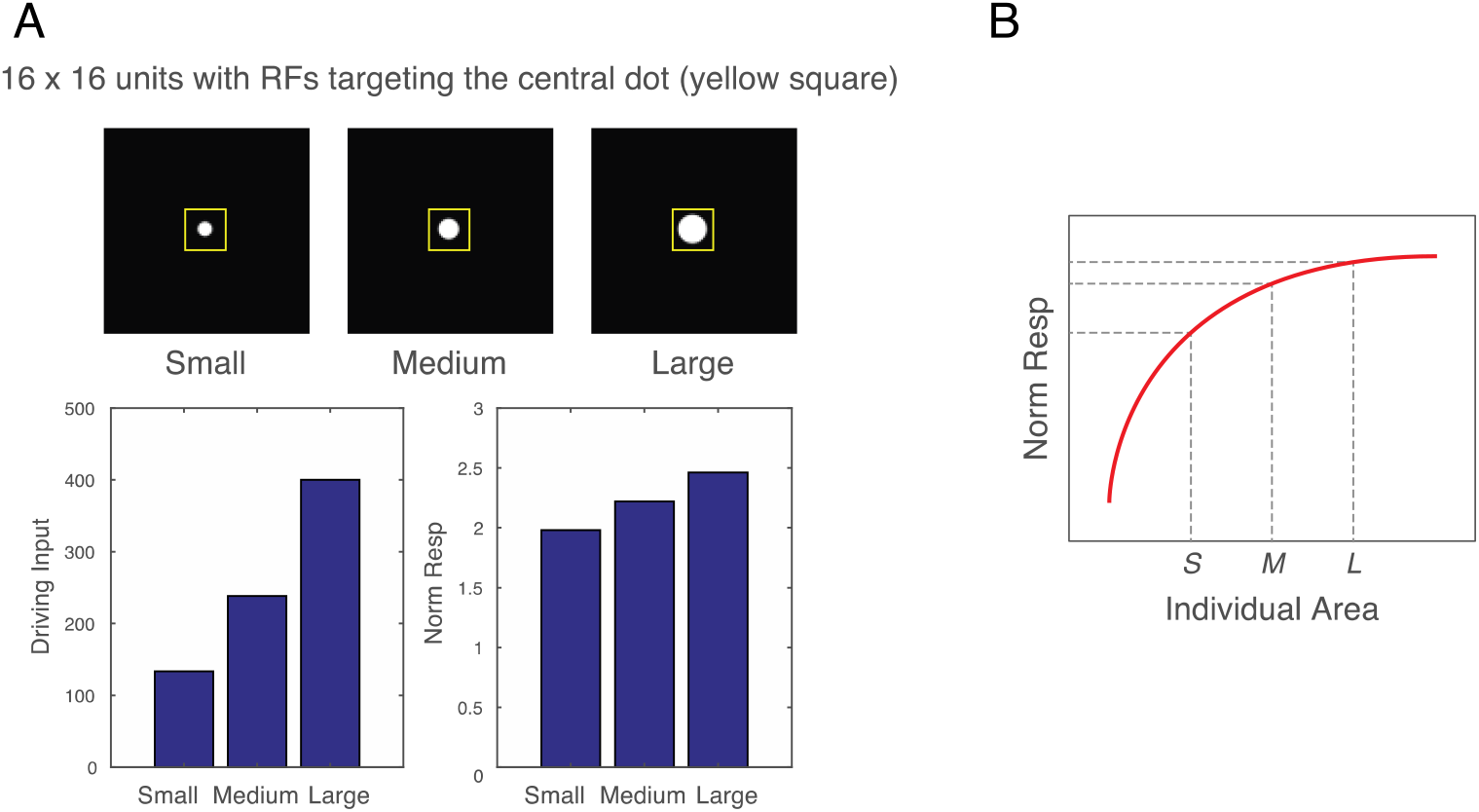
Effects of single dots. A. Images of small (radius = 3.5), medium (radius = 5), and large (radius = 7) singly presented dots were fed into the computational model, and the driving input and the normalized response of the units with the receptive fields (RF) targeting the dots were computed. As expected, driving input was nearly perfectly correlated with area of the dots (r = 0.9983). In contrast, normalized response showed a linear relationship with the radius, which meant a logarithmic relationship with area. This occurs because a larger dot involves a greater number of filters overlapping with the dot (i.e., greater driving input), but this greater number of filters leads to a greater normalization factor (increase in the denominator of divisive normalization). In other words, the normalized response becomes tempered in a non-linear way, producing a saturating normalized response as a function of increasing dot area. B. Schematic illustration of the saturating effect of normalized response for a single dot (within a hypothetical dot array) as a function of the area of the dot. Heterogeneous arrays are created by holding the total area and numerosity constant while changing individual dot size. Therefore, when medium-sized dots (M) are replaced with large dots (L), the same number of replacements must be done to go from medium-size dots (M) to small dots (S). However, because of the saturating effect, there is a greater decrease in normalized response than an increase in normalized response. Thus, the overall normalized response becomes necessarily smaller in a heterogeneous array compared to a homogeneous array.

## Notes

### Competing Interest Statement

The authors have declared no competing interest.

